# Functional screening of lysosomal storage disorder genes identifies modifiers of alpha-synuclein mediated neurodegeneration

**DOI:** 10.1101/2022.07.23.501240

**Authors:** Meigen Yu, Hui Ye, Ruth B. De-Paula, Carl Grant Mangleburg, Timothy Wu, Yarong Li, Duc Duong, Genevera I. Allen, Nicholas T. Seyfried, Ismael Al-Ramahi, Juan Botas, Joshua M. Shulman

**Affiliations:** Department of Neuroscience, Baylor College of Medicine, Houston, TX 77030, USA; Department of Neurology, Baylor College of Medicine, Houston, TX 77030, USA; Quantitative and Computational Biology Program, Baylor College of Medicine, Houston, TX 77030, USA; Department of Molecular and Human Genetics, Baylor College of Medicine, Houston, TX 77030, USA; Medical Scientist Training Program, Baylor College of Medicine, Houston, TX, 77030, USA; Departments of Biochemistry and Neurology, Emory University School of Medicine, Atlanta, GA 30322, USA; Departments of Electrical and Computer Engineering, Computer Science, and Statistics, Rice University, Houston, TX, 77030, USA; Jan and Dan Duncan Neurological Research Institute, Texas Children’s Hospital, Houston, TX, USA; Center for Alzheimer’s and Neurodegenerative Diseases, Baylor College of Medicine, Houston, TX, 77030, USA

**Keywords:** Parkinson’s disease, Gaucher disease, cholesterol, lysosome, *Drosophila*, pleiotropy, proteomics, sphingolipid, ganglioside

## Abstract

Heterozygous variants in the *glucocerebrosidase* (*GBA*) gene are common and potent risk factors for Parkinson’s disease (PD). *GBA* also causes the autosomal recessive lysosomal storage disorder (LSD), Gaucher disease, and emerging evidence from human genetics implicates many other LSD genes in PD susceptibility. We have systemically tested 88 conserved fly homologs of 38 human LSD genes for requirements in the aging adult *Drosophila* brain and for potential genetic interactions with neurodegeneration caused by α-synuclein (αSyn), which forms Lewy body pathology in PD. Our screen identifies 15 genetic enhancers of αSyn-induced progressive locomotor dysfunction, following knockdown of fly homologs of *GBA* and other LSD genes with independent support as PD susceptibility factors from human genetics (*SCARB2, SMPD1, CTSD, GNPTAB, SLC17A5*). For several genes, results from multiple alleles support dose-sensitivity and context-dependent pleiotropy in the presence or absence of αSyn. Homologs of 2 genes causing cholesterol storage disorders, *Npc1a / NPC1* and *Lip4 / LIPA*, were independently confirmed as loss-of-function enhancers of αSyn-induced brain structural degeneration. The enzymes encoded by several modifier genes are upregulated in αSyn transgenic flies, based on unbiased proteomics, revealing a possible compensatory response. Overall, our results reinforce the important role of lysosomal genes in brain health and PD pathogenesis, and implicate several metabolic pathways, including cholesterol homeostasis, in αSyn-mediated neurodegeneration.

## INTRODUCTION

Parkinson’s disease (PD) is a common and incurable neurodegenerative disorder with strong evidence for genetic etiology^1,2^. Heterozygous carriers for variants in the *glucocerebrosidase* (*GBA*) gene have an approximately 5-fold increased risk of PD, and *GBA* variants also modify PD clinical manifestations, causing more rapid progression and susceptibility for dementia^3^. Whereas partial loss-of-function is associated with PD, complete or near-complete loss of *GBA* causes Gaucher disease, a recessive lysosomal storage disorder (LSD)^4,5^. *GBA* encodes the lysosomal enzyme glucocerebrosidase (GCase), which catalyzes the breakdown of glucosylceramide, a substrate that accumulates along with other, more complex sphingolipids in Gaucher disease and possibly PD^6^.

There are more than 50 different LSDs, which are similarly characterized by defects in lysosomal biogenesis and/or function, and lead to heterogeneous clinical manifestations, including neurodegeneration in many cases^7^. Emerging evidence from human genetics suggests that, beyond *GBA*, other LSD genes, may also influence PD susceptibility. For example, heterozygous carriers of loss-of-function variants in *SMPD1*, which cause Niemann Pick Disease type A/B, have been shown to increase PD risk ^8^. Moreover, in independent studies, an aggregate burden of rare, damaging variants was associated with PD, and this relation was robust to exclusion of *GBA*^9,10^. Although the rarity of LSD gene variants limits statistical power to definitively establish the responsible genes, suggestive evidence implicates possible roles for *CTSD, SLC17A5*, and *ASAH1*. Lastly, based on genome-wide association study meta-analysis, common variants implicate several other LSD genes at PD risk loci, including *SCARB2, GRN, GUSB, GALC*, and *NAGLU*^11^.

While the precise mechanism by which *GBA* variants affect PD risk remain unknown, substantial evidence points to interactions with α-synuclein protein (αSyn), which aggregates to form Lewy body pathology. αSyn disrupts endolysosomal trafficking, including transport of GCase and other lysosomal enzymes, leading to reduced enzymatic activity and metabolic perturbations^12,13^. Reciprocally, loss of GCase may promote Lewy body pathology due to increased αSyn protein and aggregation^14^, resulting from impaired lysosomal autophagy^15^ and sphingolipid substrate accumulation^13,16,17^. In this study, we leverage a versatile *Drosophila* model to systematically test the hypothesis that other LSD genes may similarly interact with α-synuclein-mediated neurodegenerative mechanisms. Our results highlight requirements for many LSD genes in the maintenance of central nervous system (CNS) structure and function, and further reinforce links with PD pathogenesis.

## RESULTS

### Associations of LSD genes with PD risk

Previously, we and others have discovered evidence for an aggregate burden of rare genetic variants among LSD genes in association with PD risk^9,10^. Although several LSD genes have also been implicated at susceptibility loci from PD genome-wide association studies^11^, to our knowledge, a systematic analysis for common variant associations across all LSD genes has not been performed. Leveraging publicly available summary statistics from 56,306 PD cases and 1.4 million control subjects^11^, we used the multi-marker analysis of genomic annotation (MAGMA)^18^ tool to examine 51 LSD genes for enrichment of variants associated with PD.

MAGMA computes an overall gene-set test statistic considering all variants falling within gene intervals, including adjustments for gene size and regional linkage disequilibrium. The full LSD gene set was significantly enriched for variants associated with PD risk (n=51 loci, p=0.0011) (Supplemental Table 1). In order to identify possible drivers for the gene set association, we examined MAGMA output considering each of the LSD genes independently. These results identify *GBA* and 9 other LSD genes with aggregate evidence for common variant associations (p < 0.05): *IDUA, SCARB2, CLN8, GNPTAB, ARSA, GALC, CLN5, NAGLU*, and *CTSD*. We also performed a sensitivity analysis showing that the LSD gene set association remains significant after excluding either *GBA* (n=50 loci, p=0.014) or the top 3 genes (*GBA* plus *SCARB2* and *IDUA*) (n=47 loci, p=0.03), which are similarly localized to regions with genome-wide significant associations in the dataset. These results reinforce the genetic connection between the genes that cause LSDs and PD risk, revealing an important role for common variant associations.

### Screen for LSD gene modifiers of α-synuclein-mediated neurodegeneration

Human genetic studies are underpowered to comprehensively resolve all of the LSD genes contributing to PD risk and pathogenesis^9^. As a complementary approach, and to systematically test the hypothesis that LSD genes may broadly interact with αSyn-mediated neurodegeneration, we implemented a cross-species strategy. Pan-neuronal expression of the human *SNCA* gene in the fruit fly, *Drosophila melanogaster*, causes Lewy body-like αSyn aggregates, lysosomal stress, dopaminergic and other neuronal loss, and progressive locomotor impairment^19,20^. We therefore performed a genetic modifier screen examining for interactions between homologs of human LSD genes and αSyn-induced neurodegeneration, using locomotor behavior as a readout for CNS function (Figure 1). Out of 54 human LSD genes, 40 (74%) are conserved in flies (Supplemental Table 2). Overall, our screen considered 88 homologs (many genes had multiple conserved homologs), and we obtained 265 distinct strains for genetic manipulation, including RNA interference (RNAi) and other available alleles (Supplemental Table 3). We employed an automated locomotor behavioral assay based on the *Drosophila* negative geotactic response, which is highly amenable for high-throughput genetic screening^21,22^.

**FIGURE 1.**
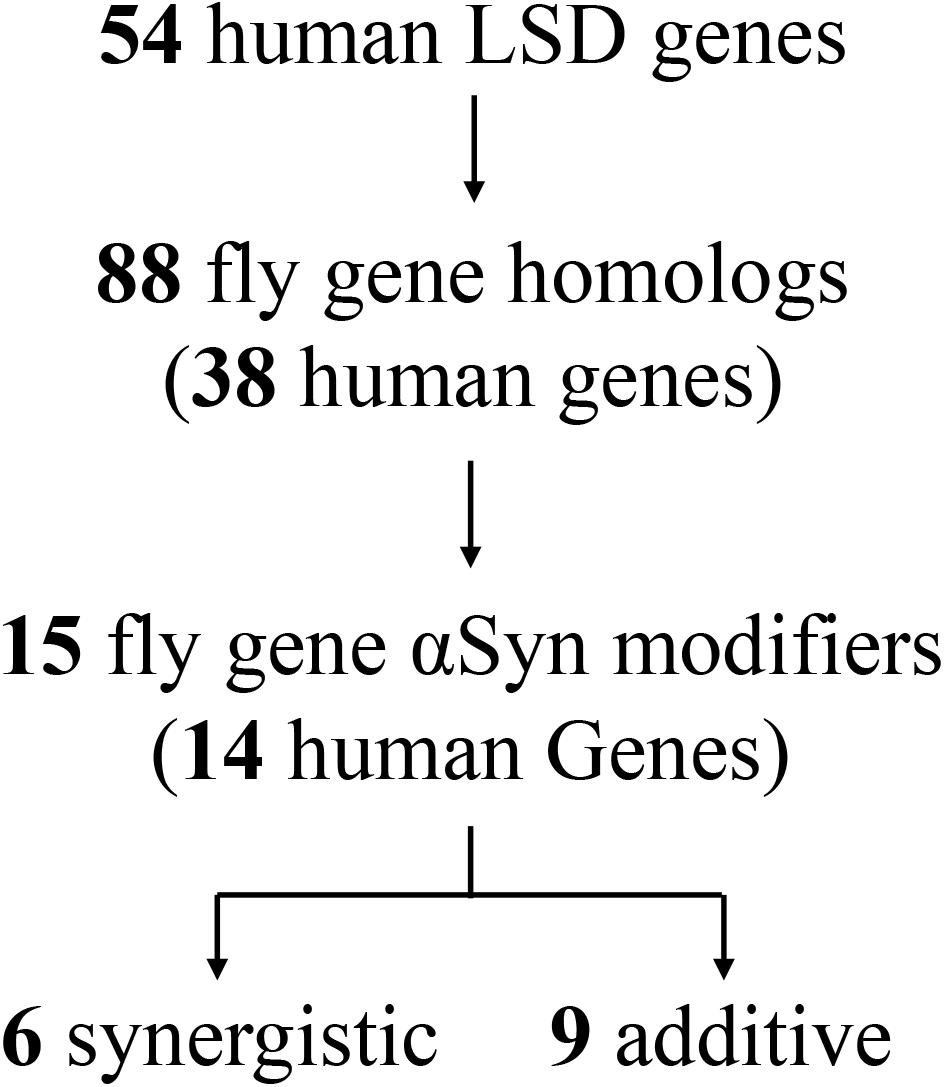
Study flowchart. Out of 54 human LSD genes, 88 are conserved in *Drosophila* and have available lines for genetic screening. 265 genetic fly strains, including RNAi and insertional or classical alleles, were tested for interaction with the locomotor phenotype induced by pan-neuronal α-synuclein (αSyn) expression. Screening revealed 15 fly genes for which gene loss significantly enhances the αSyn phenotype. Based on further tests of all modifying strains in the absence of αSyn, genes were further classified as synergistic or additive. Synergistic genes were defined as having *at least one* modifying allele in which there was no significant evidence of locomotor impairment in the absence of αSyn.

Compared to controls (*elav-GAL4 / +*), pan-neuronal expression of human αSyn (*elav> αSyn*) causes progressive locomotor impairment (Figure 2A). RNAi transgenes targeting the fly homologs of human LSD genes were coexpressed throughout the nervous system with *αSyn* (*elav>αSyn + elav>RNAi*), and climbing speed of adult flies was evaluated longitudinally between 1 and 3 weeks of age. *Drosophila* RNAi transgenic lines are designed for optimal specificity^23,24^. To further minimize the possibility of off-target effects, we only considered genes as modifiers when supported by consistent evidence from at least 2 independent RNAi strains or other alleles. Overall, our screen identified 15 fly genetic modifiers of αSyn, homologous to 14 human LSD genes (2 paralogs of *SCARB2* were identified as modifiers) (Table 1). In all cases, genetic manipulations predicted to reduce the function of LSD gene homologs (RNAi knockdown or other loss-of-function alleles) enhanced the *elav>αSyn* locomotor phenotype (Figure 2B).

**FIGURE 2.**
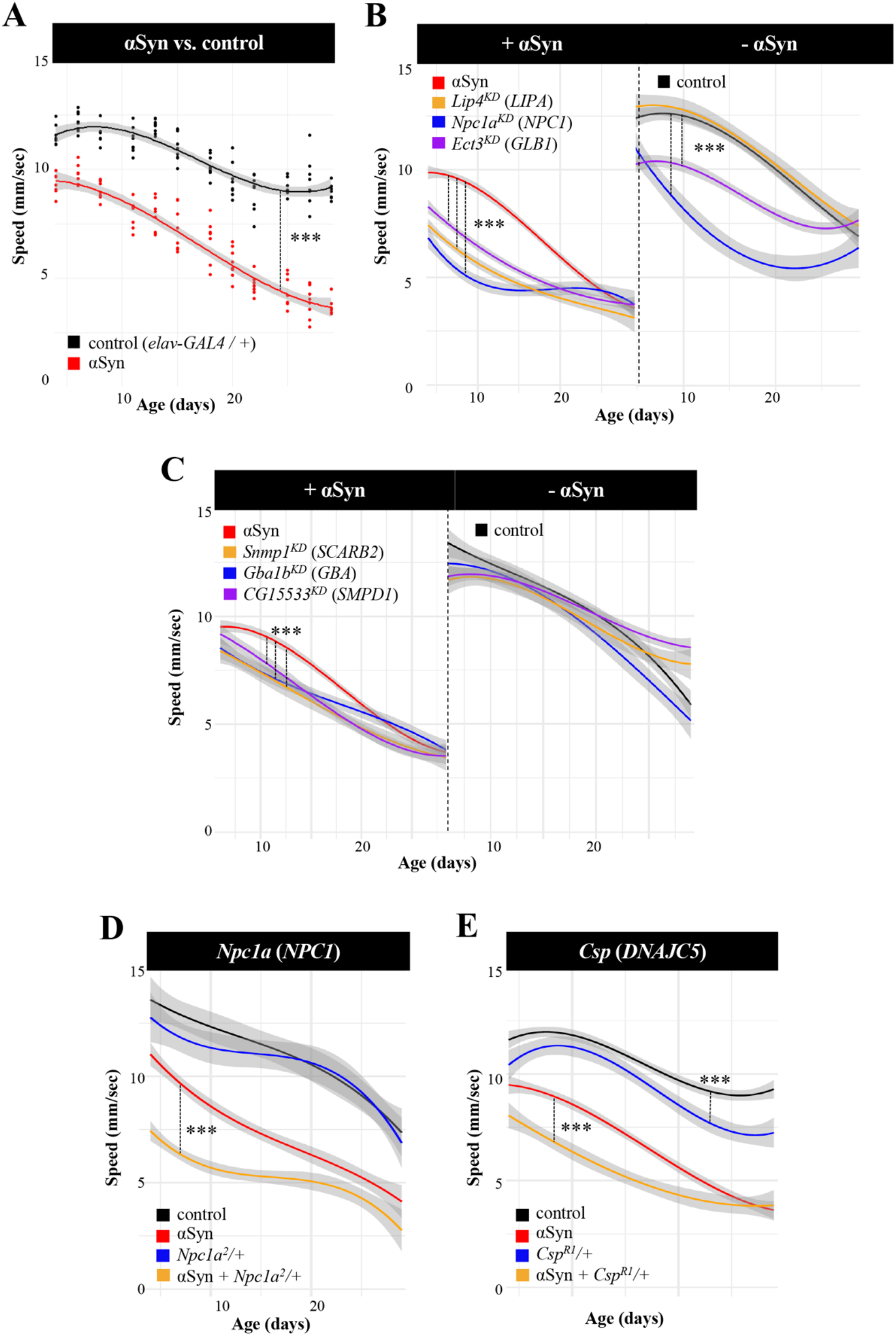
Lysosomal storage disorder (LSD) gene homologs show dose-sensitivity and context-dependent pleiotropy in *Drosophila*. (A) Pan-neuronal expression of human α-synuclein (*elav > αSyn*) induces progressive locomotor impairment. (B) Pan-neuronal knockdown (KD) of LSD gene homologs with RNA-interference (RNAi) enhances the αSyn locomotor phenotype. In the absence of αSyn, KD (*elav >* RNAi) causes no (*LIPA/Lip4*: *v31021*), moderate (*GLB1/Ect3: 3132R1*), or severe (*NPC1/Npc1a: v105405*) toxicity. (C) Additional synergistic gene modifiers enhance αSyn following KD, but do not cause significant locomotor impairment in the absence of αSyn (*SCARB2/Snmp1: v42496; GBA/Gba1b: v21336;* and *SMPD1/CG15533: v42520*). (D and E) Heterozygous loss-of-function alleles of *Npc1a* (C) and *Csp* (D) dominantly enhance αSyn. Climbing speed was assessed longitudinally including at least 11 aged time points over 30 days (n > 6 replicates of 15 animals each). ***, p<5×10^−5^. See also Supplemental Figure 1.

**TABLE 1.** Lysosomal storage disorder (LSD) gene modifiers of α-synuclein (αSyn). Human LSD genes and *Drosophila* homologs identified as modifiers are indicated. Human genes with additional supportive evidence as PD risk loci from human genetics are noted with an asterisk (*). Based on further tests to establish αSyn-dependent or independent activity, genes were further classified as synergistic (S) or additive (A). Synergistic genes were defined as having at least one modifying allele in which there was no significant evidence of locomotor impairment in the absence of αSyn. Additive genes were characterized by alleles that consistently produced locomotor phenotypes in the absence of αSyn. All modifier genes had at least one allele establishing αSyn-independent functional requirements in the fly nervous system.

|  | Human Gene | Fly Gene | Interaction |
| --- | --- | --- | --- |
| Glycosaminoglycan | *GLB1* | *Ect3* | A |
|  | *IDS* | *Ids* | S |
|  | *IDUA* * | *Idua* | A |
|  | *MAN2B1* | *LManII* | A |
| | *MANBA* | β-*Man* | A |
| Sphingolipid | *GBA* * | *Gba1b* | S |
|  | *SCARB2* * | *Dsb* | A |
|  |  | *Snmp1* | S |
|  | *SMPD1* * | *CG15533* | S |
| Chol. | *LIPA* | *Lip4* | S |
|  | *NPC1* | *Npc1a* | S |
| Other | *CTSD* * | *CG10104* | A |
|  | *DNAJC5* | *Csp* | A |
|  | *GNPTAB* * | *Gnptab* | A |
|  | *SLC17A5* * | *MFS3* | A |

Since LSDs are frequently associated with neurologic manifestations, we reasoned that knockdown of *Drosophila* homologs would likely cause CNS dysfunction in many cases, independent of αSyn expression. Therefore, we also tested all modifying lines from our screen (35 RNAi and other alleles) to examine the consequences for locomotor behavior in the absence of αSyn (e.g., *elav>RNAi* versus *elav-GAL4 / +*). These experiments revealed a range of phenotypic severity (Figure 2B and Supplemental Figure 1), with some RNAi lines showing little or no locomotor phenotype (e.g., *LIP4 / Lip4*) and others with mild (e.g., *GLB1 / Ect3*) or more substantial, age-dependent impairments (e.g., *NPC1 / Npc1a*). Based on these results, we classified the 15 genes as either “additive” or “synergistic” modifiers of αSyn (Table 1). Synergistic modifiers had at least one modifying allele in which there was no significant evidence of locomotor impairment in the absence of αSyn (Figure 2C). Overall, 6 of 15 genes, including *Gba1b*, the *Drosophila* homolog of *GBA*, showed evidence of synergistic interactions with αSyn mediated neurodegeneration. The remaining “additive” genetic modifiers were characterized by alleles that consistently produced locomotor phenotypes in the absence of αSyn. Notably, all LSD modifier genes had at least one allele tested revealing αSyn-independent functional requirements in the aging nervous system.

In humans, *GBA* reveals dose-sensitive and pleiotropic effects on disease traits, with near complete loss-of-function causing Gaucher disease and partial loss increasing susceptibility for PD. In *Drosophila*, our screen results also suggest that many LSD gene homologs may have similar dose-dependent relationships (Supplemental Figure 1). For 3 loci, including *GBA* (*Gba1b*), *IDS* (*Ids*), and *LIPA* (*Lip4*), knockdown with multiple, independent RNAi transgenes targeting each of these genes showed consistent enhancement of αSyn-induced locomotor impairment, but differential requirements in the absence of αSyn. For selected genes, classical mutant alleles were also available and permitted evaluation of potential heterozygous interactions. Strikingly, heterozygous loss-of-function alleles for both *Npc1a* and *Csp*, homologous to human *NPC1* and *DNAJC5*, respectively, dominantly enhanced αSyn, but caused little to no phenotype when examined on their own (Figure 2D, E and Supplemental Figure 1). By contrast, RNAi-knockdown of both genes induced a marked locomotor phenotype independent of αSyn (Figure 2B and Supplemental Figure 1). Overall, these results are consistent with a model in which partial loss of function for multiple LSD genes may enhance αSyn neuropathology, as with *GBA*-PD, but that more complete loss of gene function may compromise CNS function, as in neuronopathic Gaucher disease and many other LSDs.

### Cholesterol metabolism and α-synuclein mediated neurodegeneration

LSDs are commonly classified based on the metabolic pathways disrupted and characteristic type of substrate accumulation. For example, *GBA, SCARB2*, and *SMPD1*, which are collectively implicated in PD risk are also jointly involved in sphingolipid metabolism; fly homologs of all 3 genes (*Gba1b, Dsb*, and *CG15533*, respectively) were identified in our screen as synergistic, loss-of-function enhancers of αSyn. Among our results, *Npc1a* and *Lip4* are both fly homologs of genes causing the human cholesterol storage disorders, Niemann Pick Disease type C (*NPC1*) and cholesterol ester storage disease (*LIPA*). Both *Npc1a* and *Lip4* were notable for robust, synergistic enhancement, consistent with dose-sensitive interactions with αSyn-mediated neuronal injury (Figure 2B, D). Whereas *Lip4* has not been well-characterized in flies, loss of *Npc1a* causes the accumulation of cholesterol, defective synaptic transmission, and ultimately, neurodegeneration, similar to human Nieman Pick type C^25^.

To further examine the interactions between αSyn and *Npc1a* and *Lip4*, we used an independent retinal neurodegeneration assay. Following expression of αSyn in adult photoreceptors using the *Rh1-GAL4* driver (*Rh1>* α*Syn*), age-dependent structural degeneration leads to mild vacuolar change at 15 days, based on hematoxylin and eosin staining of tangential sections through the retina (Figure 3)^26^. RNAi-mediated *Lip4* knockdown or a heterozygous *Npc1a* loss-of-function allele significantly increased αSyn-induced retinal degeneration, but similar changes were not seen in corresponding controls. These results are consistent with the findings from our screen and further implicate lysosomal regulators of cholesterol metabolism in αSyn-mediated neurodegeneration.

**FIGURE 3.**
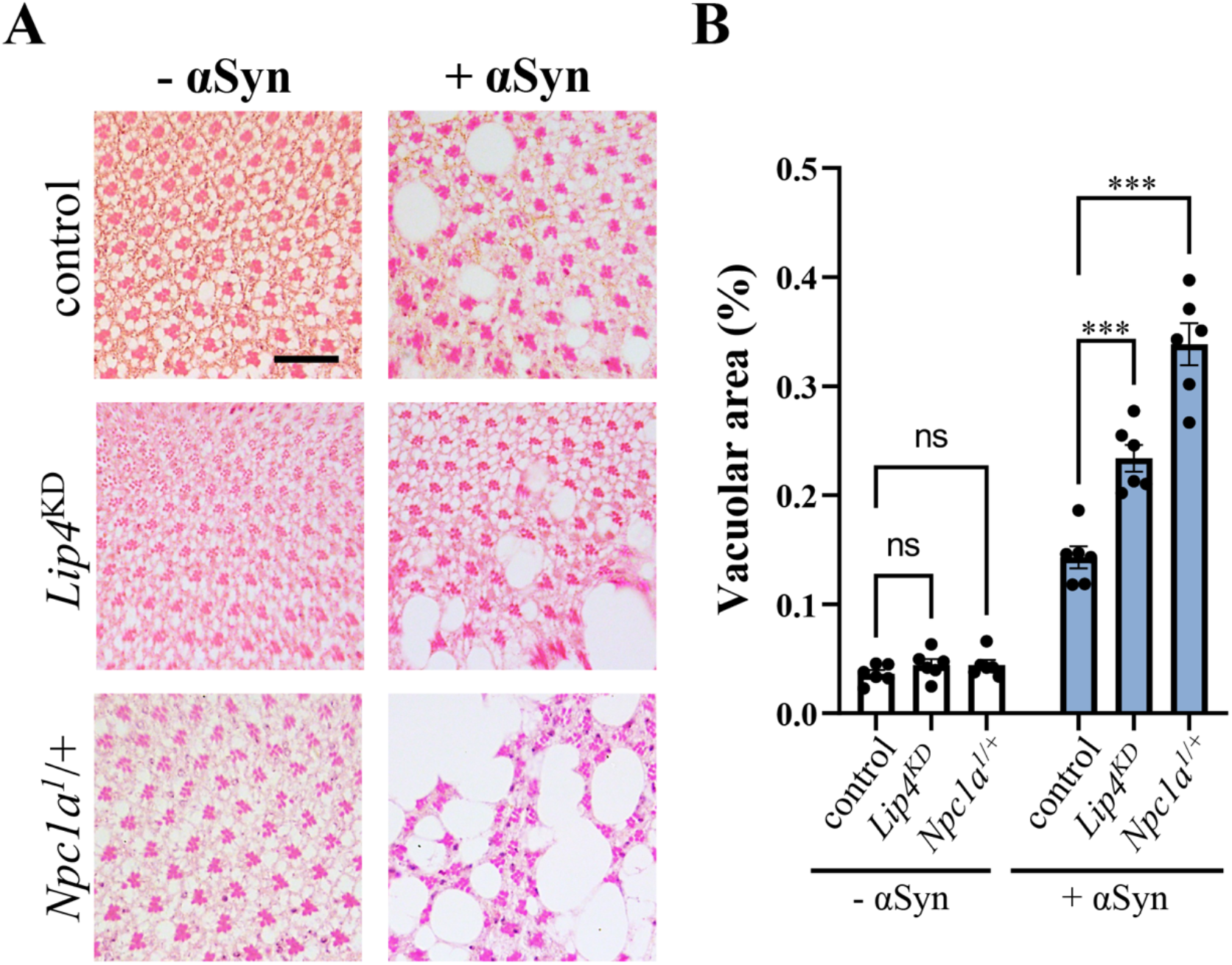
Cholesterol storage disorder gene homologs enhance α-synuclein (αSyn) induced retinal degeneration. (A) Expression of αSyn in the adult retina (*Rh1>αSyn*) causes progressive neurodegeneration compared with controls (*Rh1-GAL4 / +*). Knockdown (KD) of *Lip4* (*v31021*) or heterozygosity for a *Npc1a* loss-of-function allele enhances αSyn-mediated tissue destruction. Tangential retinal sections from 15 day-old animals were stained with hematoxylin and eosin. (B) Quantification based on extent of vacuolar changes (vacuole area / total area) from at least n=6 animals per genotype. Statistical comparisons were made using unpaired t-tests, followed by Dunnett’s post-hoc test. Error bars represent the standard error of the mean. See also Supplemental Figure 3. ***, p<0.001; ns, not significant. Scale bar=20*μ*m. See also Supplemental Figure 2 for results using additional RNAi and alleles.

### α-synuclein pathology causes altered expression of LSD enzymes

In order to further explore underlying molecular mechanisms, we generated unbiased mass-spectrometry proteomics from αSyn (*elav>αSyn*) and control (*elav-GAL4 / +*) animals, using the same genotypes as in our locomotor screen and choosing a timepoint (10-days) that is predicted to be early in the overall pathologic progression. From these data, 49 fly homologs of human LSD proteins were detected. Strikingly, 23 *Drosophila* proteins (homologous to 17 human proteins encoded by LSD genes) were significantly differentially expressed following pan-neuronal expression of αSyn in the adult brain (Figure 4 and Supplemental Table 4), including 15 up- and 8 down-regulated proteins. Importantly, 7 proteins overlapped with genetic modifiers identified in our screen, including Npc1a as well as several proteins homologous to human LSD gene products causing mucopolysaccharidoses: GLB1/Ect3, IDUA/Idua, MAN2B1/LManII, and MANBA/Beta-Man. Collectively, these enzymes participate in the breakdown of glycosaminoglycans (Table 1). Interestingly, 6 out of 7 of the overlapping modifiers are up-regulated proteins. Since gene loss-of-function enhances the locomotor phenotype, the increased protein expression is consistent with a potential compensatory response to αSyn-mediated lysosomal stress.

**FIGURE 4.**
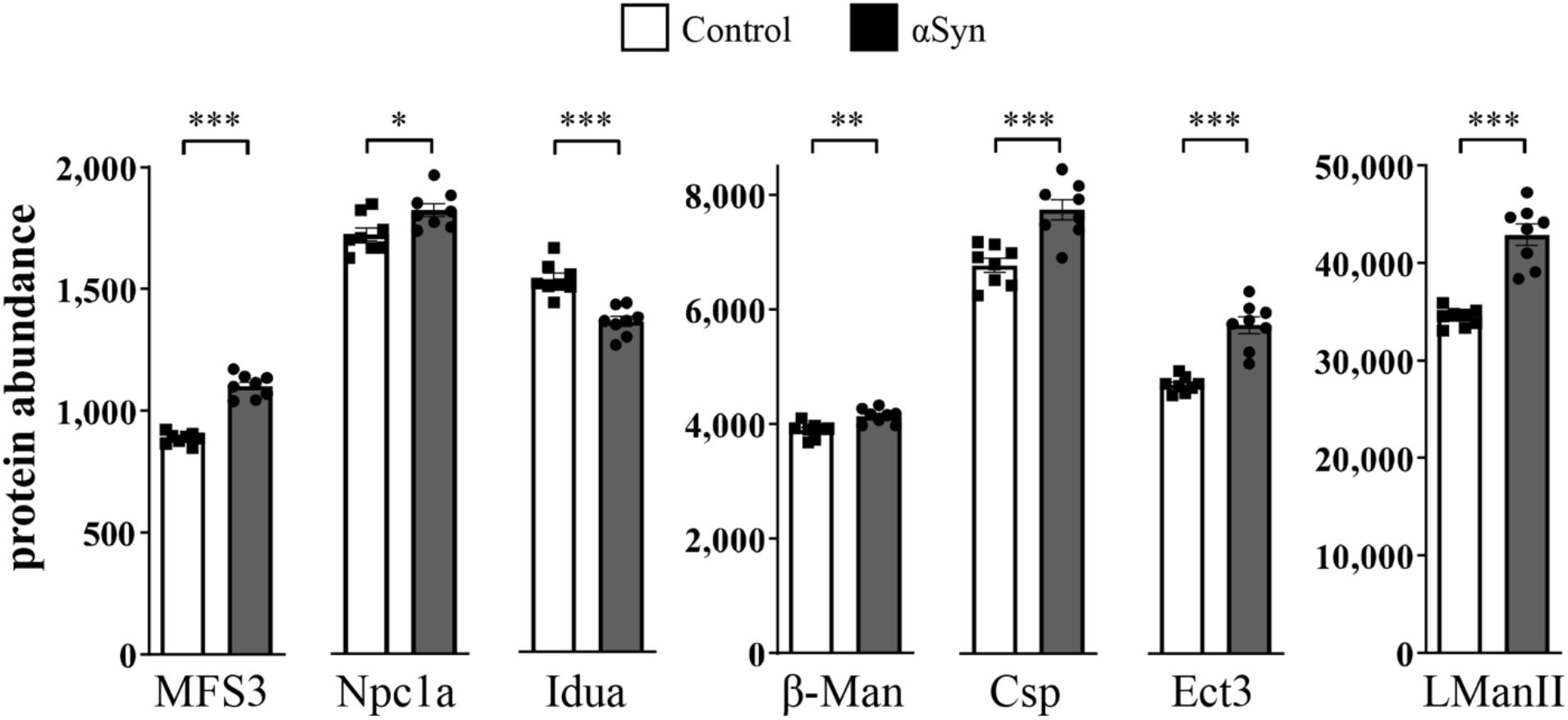
Differential expression of LSD protein homologs in flies. Comparisons of protein abundance (normalized) from fly heads are shown for *elav>αSyn* (Gray) or control (White, *elav-GAL4 / +*), based on Tandem Mass Tag proteomics. t-tests were performed for comparisons of mean abundance considering n=8 replicate samples for each genotype. Error bars represent the standard error of the mean. *, p<0.05; **, p<0.01; ***, p<0.001. All proteins shown were also significantly differentially expressed based on analyses in DESeq2 (Wald test) and following adjustment using the Benjamini-Hochberg procedure (p_adj_ < 0.05). See also Supplemental Table 4.

## DISCUSSION

Mounting evidence supports an important connection between the genetic mechanisms of LSDs and PD. Using a cross-species, functional screening strategy, we discover 14 conserved human LSD genes with homologs that robustly enhance αSyn-mediated neurodegeneration when their activity is reduced in *Drosophila* models. The majority of these genes also show evidence for αSyn-independent requirements, and in several cases our data is consistent with dose-sensitivity and context-dependent pleiotropy similar to *GBA* in PD and Gaucher disease. Two of the genes identified by our screen, *GBA* and *SMPD1*, are established PD risk genes^3,8,27^, and our results confirm and extend data from other animal and cellular models (discussed below). In other cases, the evidence from human genetics may be more modest, but our discovery of genetic interactions with αSyn mechanisms increases the possibility that these genes may be *bona fide* PD risk factors. For example, in our prior analysis of exome sequencing data, *CTSD* and *SLC17A5* showed suggestive associations with PD risk^9^; however, the rarity of variants and available sample sizes have limited statistical power to confirm such loci. Our results also provide experimental support for LSD genes, such as *SCARB2* and *GNPTAB*, which are candidates at susceptibility loci from PD GWAS^11^. Indeed, GWAS rarely identify responsible genes definitively, but instead highlight regions that usually contain many viable candidates. As a group, we and others previously showed that genes causing LSDs harbor an aggregate burden of damaging rare variants among PD cases^9,10^. Here, using available GWAS data, we also demonstrate similar significant enrichment for more common variants associated with PD risk. Overall, the LSD genes prioritized by our functional screening strategy are outstanding candidates for further investigation as PD risk factors, including using both human genetics and experimental approaches.

Animal and cellular model experimental studies highlight several plausible mechanisms for how LSD genes might interact with and enhance αSyn-mediated neurodegeneration. First, LSD gene loss-of-function may impair turnover and thereby increase accumulation of toxic αSyn species. For example, *CTSD* encodes a lysosomal cathepsin, which can directly degrade αSyn^28,29^. More indirectly, reduced LSD gene activity may promote the accumulation of undigested substrates that secondarily accelerate αSyn misfolding or aggregation. In a variety of systems, glucosylceramide and its derivatives (e.g., glucosylsphingosine) have been shown to interact with αSyn in this manner^16,17^. LSD gene dysfunction may also promote lysosomal stress and reduced autophagic flux, thereby compromising αSyn protein turnover^30,31^. Experimental manipulations of *GBA, SMPD1*, and *SCARB2—*which similarly cause sphingolipid storage disorders—are known to induce accumulation and toxicity of αSyn^13,27,32^. Beyond sphingolipid/ceramide metabolism, our screen also identifies many fly homologs of genes causing human mucopolysaccharidoses (e.g., *GLB1, IDS, IDUA, MAN2B1, MANBA*). These disorders are characterized by accumulation of glycosaminoglycans, which are long chains of repeating, negatively charged disaccharide units that can be linked to protein cores to form proteoglycans. Though perhaps less well studied than sphingolipids, recent studies suggest that glycosaminoglycan metabolites have the potential to similarly influence αSyn aggregation^33^. In addition, proteoglycans are abundant at the cell surface and are a major constituent of the extracellular matrix, having recently been implicated in the propagation and/or internalization of pathologic αSyn species^34^. Interestingly, in one recent human genetic analysis, the genes causing mucopolysaccharidoses comprised a major driver for the rare variant burden association with PD risk among LSD genes^10^. Our screen also identifies homologs of 2 causes of cholesterol storage disorders, *LIPA* and *NPC1*. Cholesterol content may also be an important modulator of interactions between αSyn and the neuronal membrane, with consequences for synaptic transmission and PD pathologic progression^35,36^. In fact, conflicting epidemiologic studies have linked hypercholesterolemia to either increased^37^ or decreased^38–40^ PD risk, whereas other studies have found no such association^41^. To date, there has been no definitive genetic evidence for association of *NPC1* with PD risk^42,43^; however, subclinical metabolic perturbations^44,45^ or parkinsonian signs^46,47^ have been reported in heterozygous carriers of loss-of-function variants. Lastly, it is possible that disparate molecular pathways interact and feedback on one another within the lysosomal milieu. For example, loss of *NPC1* in Niemann Pick Disease type C leads to accumulation not only of cholesterol, but also secondary perturbations in sphingolipids and glycosaminoglycans^48^. Thus, it is conceivable that subtle perturbations affecting one enzymatic transformation might trigger a cascading metabolic failure that ultimately amplifies αSyn mediated neurodegeneration.

Ultimately, a successful mechanistic model must account for both the pleiotropic potential of LSD genes and their exquisite, dose-dependent interactions with αSyn-mediated PD mechanisms. Strikingly, whereas only a modest reduction of GCase enzymatic activity confers increased PD risk (e.g., ∼75% residual function in carriers of the *GBA*^*E356K*^ allele)^49^, Gaucher disease requires near complete loss of function (e.g., less than 15% residual GCase activity)^50^. Based on our screen, nearly all fly homologs of human LSD genes were capable of causing locomotor dysfunction independent of αSyn. This result is consistent with an obligate role for most LSD genes in the maintenance of CNS structure and function. By contrast, more modest loss-of-function in many LSD genes strongly enhanced the αSyn locomotor phenotype but caused little or no CNS dysfunction independent of αSyn. This was best exemplified by fly homologs of *NPC1* and *DNAJC5* (*Npc1a* and *Csp*, respectively) which both showed dominant, heterozygous enhancement of αSyn neurotoxicity, supporting dose-sensitive interactions. For many other genes, including the *Drosophila GBA* homolog, *Gba1b*, we also identified additional RNAi strains that support a synergistic interaction model. Prior studies using human αSyn transgenic flies have differed on potential synergistic interactions following *Gba1b* manipulation, but these studies have relied on either different alleles and tissue-specific drivers or distinct phenotypic assays^51,52^. Dose-sensitive interactions may arise from positive feedback between aging, αSyn toxicity, and progressive lysosomal dysfunction. Data from multiple experimental models highlight how αSyn potently disrupts endosomal trafficking pathways, including delivery of GCase and other hydrolases to the lysosome^12,13^, and thus may potentiate the impact of partial genetic loss-of-function in LSD genes. These pathologic interactions may be further amplified by aging, which among myriad cellular changes, is accompanied by reduced lysosomal proteostasis (autophagy)^53^. The substantial impact of αSyn on the lysosome is reflected in our *Drosophila* proteomic profiles, revealing perturbations in the expression of numerous LSD proteins. In particular, the up-regulation detected for 6 out of 7 modifier genes is consistent with a possible compensatory cellular response, since genetic manipulation in the opposite direction (RNAi knockdown) worsened αSyn-induced locomotor impairment.

The strengths of this study include the cross-species strategy, systematic consideration of multiple alleles for all LSD gene targets, longitudinal data collection, and an analytic framework that accounts for the potential impact of aging. The high-throughput locomotor screening assay is sensitive to early consequences of CNS dysfunction that precede cell death and subsequent structural degenerative changes. One important potential limitation is that all genetic manipulations with RNAi were targeted exclusively to neurons. Many LSD genes, including *GBA*, are also expressed in glia, where they may also have important roles in the maintenance of CNS structure/function^54^. While we also tested classical mutant alleles, which are predicted to affect gene function globally, these reagents were only available for a subset of LSD gene homologs. Lastly, a minority of LSD genes are non-conserved in the *Drosophila* genome and were therefore not examined, including some with evidence for association with PD risk from human genetics (e.g., *ASAH1, GUSB*). Nevertheless, our results confirm and extend the strong genetic link between LSDs and PD and highlight many promising genes and metabolic pathways for further study in PD risk, pathogenesis, and therapy.

## MATERIALS AND METHODS

### LSD gene set enrichment analysis from PD GWAS

PD GWAS summary statistics^11^ were analyzed using MAGMA v1.10^18^. Gene location and European reference files for the GRCh37 genome build were downloaded from MAGMA webpage (https://ctg.cncr.nl/software/magma), and the BEDTools v2.26.0^55^ *intersect* function was used to interrogate SNPs from the PD GWAS summary statistics. MAGMA annotation, gene analysis and gene-set analysis steps were performed using default parameters. The list of 54 LSD genes^9^ (Supplemental Table 1) was used under the *--set-anot* parameter for the gene-set analysis, and selected genes were excluded for the sensitivity analysis. Three X-linked genes (*GLA, IDS*, and *LAMP2*) were not considered since variants were not available.

### Fly stocks and husbandry

Human *α-synuclein* transgenic lines with codon optimization for *Drosophila* were previously described ^20^. For locomotor screening, a recombinant second chromosome line harboring 2 *UAS-α-synuclein* insertions was used, as in prior studies^22^. For retinal histology, a third chromosome insertion was employed, also from previous work^26^. The *GAL4-UAS* system^56^ was used for ectopic expression of both αSyn and RNA-interference (RNAi) transgenes. For pan-neuronal expression, we used *elav*^*c155*^*-GAL4* (ref. 57), which is available from the Bloomington *Drosophila* Stock Center (BDSC; Bloomington, IN, USA). For expression in retinal photoreceptors, we used *Rh1-GAL4* (second chromosome insertion)^26,58^. For the locomotor screen of conserved LSD genes, virgin females obtained from RNAi or other allelic strains were crossed to male *elav*^*c155*^*-GAL4 / Y; UAS-SNCA / Cyo,GAL80*. To evaluate αSyn-independent effects, RNAi or other genetic strains, potential modifier strains were crossed with *elav*^*c155*^*-GAL4 / Y* or *w*^*1118*^ */ Y* males. All modifier gene manipulations (RNAi or alleles) were tested in heterozygosity. Crosses and progeny were established and maintained at 25°C. For secondary modifier tests using the αSyn retinal degeneration assay, all crosses were established at 18°C, and progeny were shifted to 25°C within 24 hours of eclosion and aged 15 days, following published protocols^26^. RNAi transgenic strains (*UAS-RNAi*) were obtained from the Vienna Drosophila Resource Center (VDRC; Vienna, Austria)^23^, from BDSC for the Harvard Transgenic RNAi Project^24^, as well as the National Institute of Genetics Fly Stock Center (NIG; Japan). All RNAi lines used for this study are detailed in Supplemental Table 3. As a control, we used the VDRC *w*^*1118*^ genetic background strain (line 60000). Additional alleles for fly homologs of LSD genes, including transposon insertions and other classical mutations, were obtained from BDSC or requested from other laboratories, including *crq*^*KO*^ (ref. 59), *dsb*^*KO*^ (ref. 60), and *Gba1b*^*KO*^ and *Gba1a,b*^*KO*^ (ref. 61).

### Genetic screen

The *Drosophila* Integrated Ortholog Prediction Tool (DIOPT)^62^ was used to identify all conserved fly homologs of human LSD genes. We required consensus from at least 3 independent bioinformatic algorithms for inclusion of a fly gene homolog (DIOPT score of 3 or greater). Where multiple paralogs met our criteria, we attempted to carry forward all such candidates. In the exceptional cases where more than 5 gene paralogs were identified as potential homologs for a given human gene, a more stringent DIOPT score cut-off was used (Supplemental Table 1). The screen was conducted in 2 stages. First, 177 RNAi strains from VDRC were obtained to knock down 88 unique homologs of human LSD genes (reagents were not available for 6 conserved genes). Locomotor behavior (see below) was assayed in female adult flies at up to 5 time points between 8 and 18 days post-eclosion. Based on the results, we subsequently obtained an additional 88 lines for further screening, including independent RNAi strains (from BDSC and NIG) and other available alleles. Lastly, for validation of screen results, all modifier genes supported by evidence from multiple independent allelic strains were reevaluated in both the presence and absence of αSyn, and locomotor behavior was assayed at 11 or more time points over 30 days of aging.

### Robot-assisted locomotor assay

The negative geotaxis climbing assay was performed using a custom robotic system (SRI International, available in the Automated Behavioral Core at the Dan and Jan Duncan Neurological Research Institute), as previously described.^21^ Negative geotaxis is elicited by “tapping” vials of flies to knock them to the bottom of custom vials. After three taps, video cameras recorded and tracked the movement of animals at a rate of 30 frames per second for 7.5 seconds. For each genotype, 6-8 replicates of 15 female animals were tested in parallel (biological replicates), and each trial was repeated five times (technical replicates). Replicates were randomly assigned to positions throughout a 96-vial plate and were blinded to users throughout the duration of experiments. Quantification was based on the average climbing speed of flies included in each biological replicate. Speed of individual flies was computationally deconvoluted from video recordings. For configuration and running of the robotic assay and video acquisition, we used the following software packages: Adept desktop, Video Savant, MatLab with Image Processing Toolkit and Statistics Toolkit, RSLogix (Rockwell Automation), and Ultraware (Rockwell Automation). Additional custom software was developed for assay control (SRI graphical user interface for controlling the machine) and analysis [FastPhenoTrack (Vision Processing Software), TrackingServer (Data Management Software), ScoringServer (Behavior Scoring Software), and Trackviewer (Visual Tracking Viewing Software)].

### Adult brain histology

Fly heads from 15-day-old animals were fixed in 8% glutaraldehyde and embedded in paraffin. Tangential retinal sections (3 μm) were cut using a Leica Microtome (RM2245) and stained with hematoxylin and eosin. Retinas from at least three animals were examined and quantified for each experimental genotype. Enhancement of αSyn-induced retinal degeneration was quantified based on the severity of retinal vacuolar changes seen in stained histologic sections. We examined representative photographs taken with a 40X objective from well-oriented, intact tangential sections at a depth in which the retina achieves maximal diameter. Using ImageJ software,^63^ we recorded the area occupied by all vacuoles with a diameter greater than 4μm and divided by the total retinal area to compute a percentage.

### Proteomic analysis of LSD genes

Tandem mass tag mass-spectrometry proteomics was performed for αSyn transgenic and control flies, using the identical genotypes as the locomotor screen (*elav>aSyn* and *elav-GAL4 / +*) and following previously published protocols^64^. Homogenates were prepared from adult fly heads aged to 10 days, including 8 biological replicate samples per model consisting of approximately 50 heads each. The full dataset includes quantitation of 6610 unique proteins and is included as supplemental data. Proteins containing missing values were excluded and protein intensity values mapping to the same flybase ID were summed. For this study, our analyses were restricted to 49 *Drosophila* proteins (Supplemental Table 4); a more comprehensive, proteome-wide analysis will be reported elsewhere. Differential protein expression was calculated with DESeq2 v1.34.0^65^ using genotype as a linear regression covariate and using the *lfcShrink* function. The DESeq2 *counts* function was implemented in order to plot normalized abundance (Figure 4).

### Statistical Analysis

MAGMA analysis was performed using default parameters. An F-test was performed to compute gene p-values. For gene set analyses, p-values are converted to Z-values, and a competitive analysis is performed, returning an overall T-test p-value. Data from our robotic *Drosophila* locomotor assay was processed by first calculating mean and standard deviation values for climbing speed across the replicates of each genotype tested using the *mean* and *sd* functions in R^66^. Each experimental genotype was compared against all control genotypes tested on the robotic assay at the same time for all downstream statistical analyses. Within-tray analyses were conducted to minimize differences between groups of flies (potential batch effects). We employed longitudinal mixed effects models in our analyses to better represent the age-dependent changes which we hypothesized were modified by differences in genotype. These models were implemented using the *lme4* package in R^67^. We used a random intercept term to model the mean climbing speed of each genotype and smoothing splines (cubic B-splines) to capture non-linear trends over time^68^. We tested the differences between all possible pairs of genotypes within each tray by testing the interactions between genotypes and their B-splines using one-way ANOVA (*aov* function in R) with three nested statistical models of increasing complexity: (i) genotype, (ii) genotype + time, and (iii) genotype*time. The genotype-only model examined mean shifts in climbing speed between genotypes without accounting for changes over time; the genotype+time model additionally considered non-linear time trends; and the genotype*time model also considered interactions leading to changes in spline slope between genotypes. We accepted p-values derived from the most complex statistical model (i, ii, or iii) meeting our significance threshold of α=5×10^−5^. For the histology assay, comparisons of vacuolar degenerative change were made using a 2-tailed student’s unpaired t-test followed by Dunnett’s post-hoc test for multiple comparisons, implemented in GraphPad Prism. Except as noted above, differential expression analysis of proteomics data was performed using default DESeq2 parameters, which implements a Wald test, and differentially expressed proteins were filtered using a significance threshold of Benjamini-Hochberg adjusted p-value < 0.05. For visualization (Figure 4), t-tests were used to compare the normalized abundance levels between control and αSyn flies for each selected gene.

## ACKNOWLEDGEMENTS

We thank our colleagues, Drs. Chun Han, Michael Hoch, and Linda Partridge for generously providing *Drosophila* strains. We also thank the Bloomington *Drosophila* stock center, the Vienna *Drosophila* RNAi Center, the TRiP at Harvard Medical School, the National Institute of Genetics, Japan, and FlyBase. We are grateful to Dr. Laurie Robak for helpful discussions and feedback on the manuscript.

