## Supplemental Figures for "Functional screening of lysosomal storage disorder genes identifies modifiers of alpha-synuclein mediated neurodegeneration"

**SUPPLEMENTAL FIGURE 1**

Data from the locomotor screen validation phase is shown for all modifiers of the α-synuclein (αSyn) locomotor phenotype. Pan-neuronal expression of human α-synuclein (Red: *elav > αSyn*) induces progressive locomotor impairment versus control flies (Green: *elav-GAL4* / +). Homologs were manipulated using RNA-interference (RNAi) or using loss-of-function alleles (allele). Each modifier gene was tested in both the presence (Purple: *elav>αSyn* + modifier) or absence (Blue: *elav*>RNAi or *elav-GAL4* + allele) of αSyn. Statistical analysis (one-way ANOVA) was performed from longitudinal mixed effects models with smoothing splines. Significance testing examined whether modifier genes enhance the αSyn-induced locomotor impairment [p(+ syn)] and whether gene manipulations cause locomotor phenotypes independent of αSyn [p(-syn)]. The 2 comparisons (*i.* and *ii.*) are indicated on the first plot shown below. We classified modifier strains based on the severity of phenotype produced independent of αSyn, including (A) no / mild, (B) moderate, or (C) severe toxicity.

**(A) no / mild toxicity independent of αSyn**

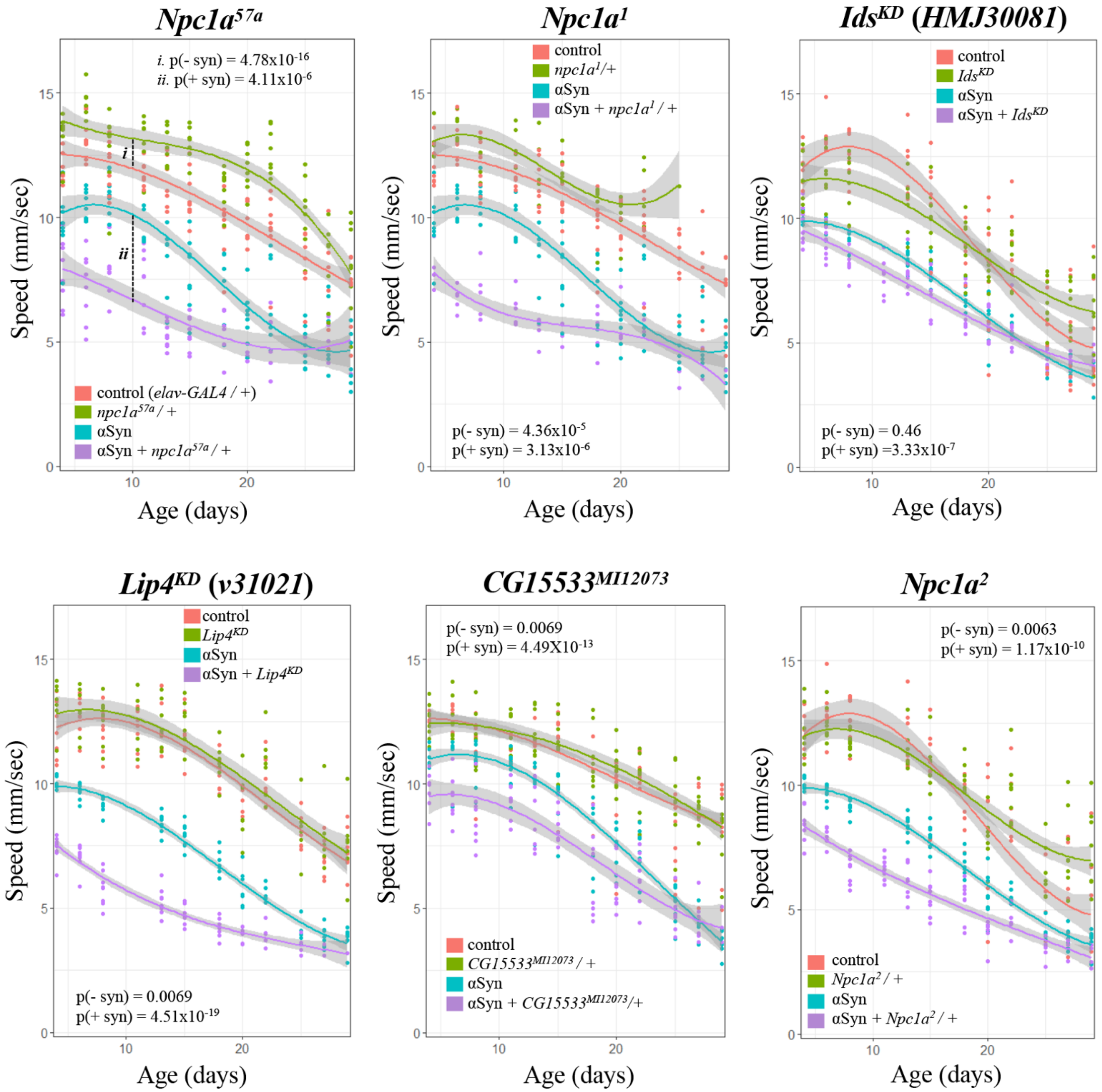

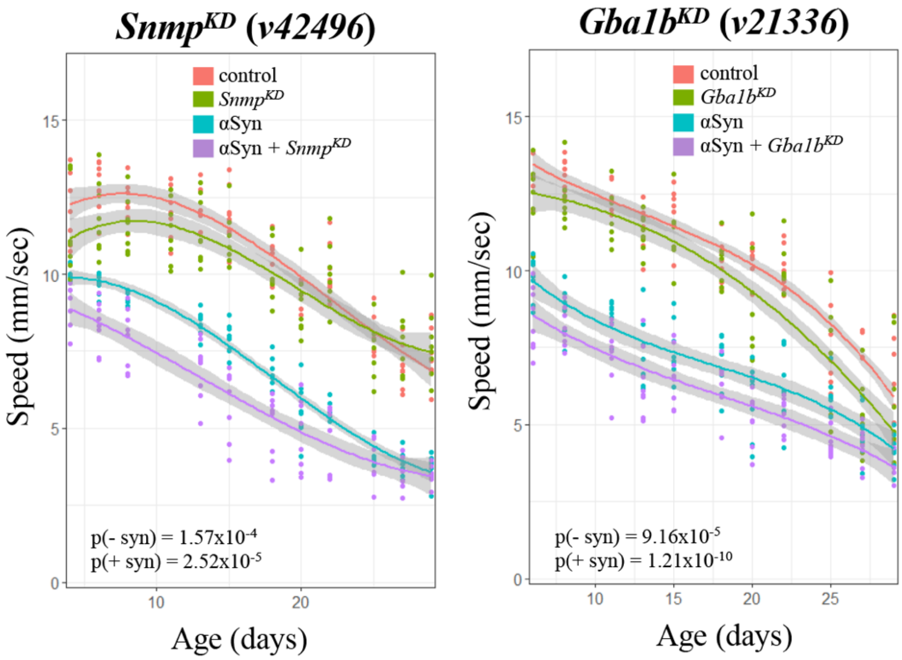


**(B) moderate toxicity independent of αSyn**

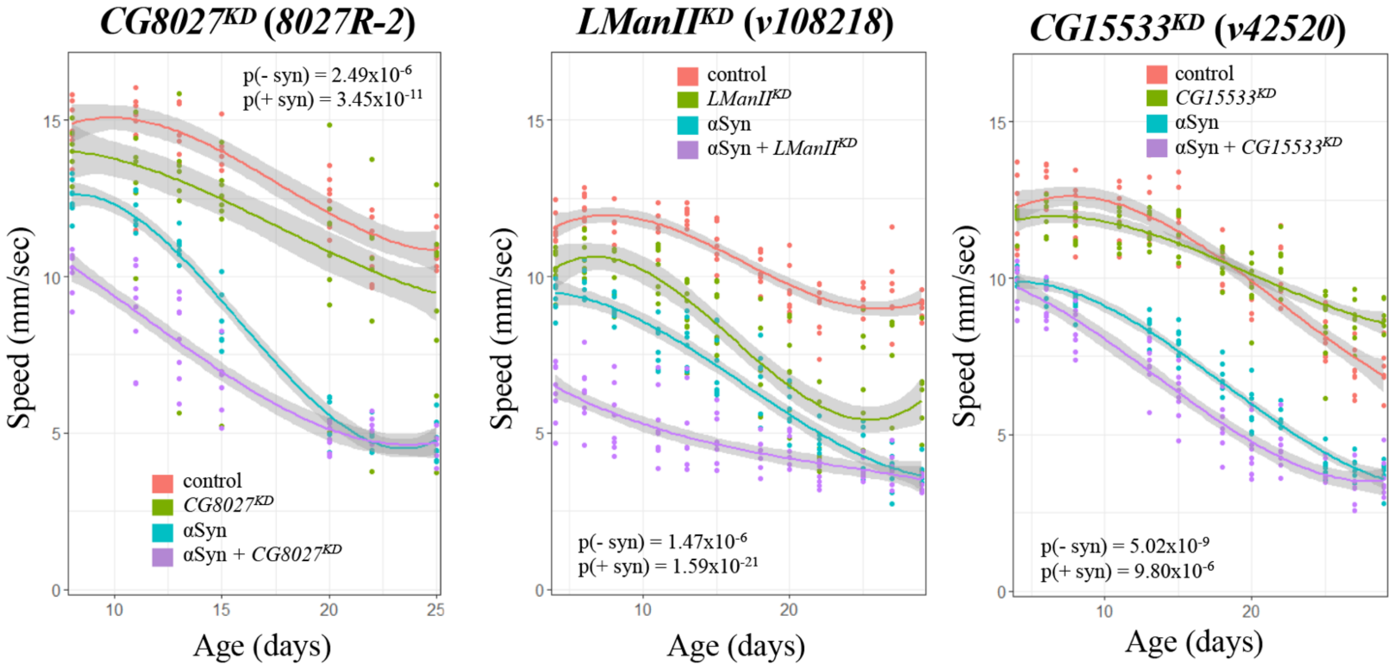

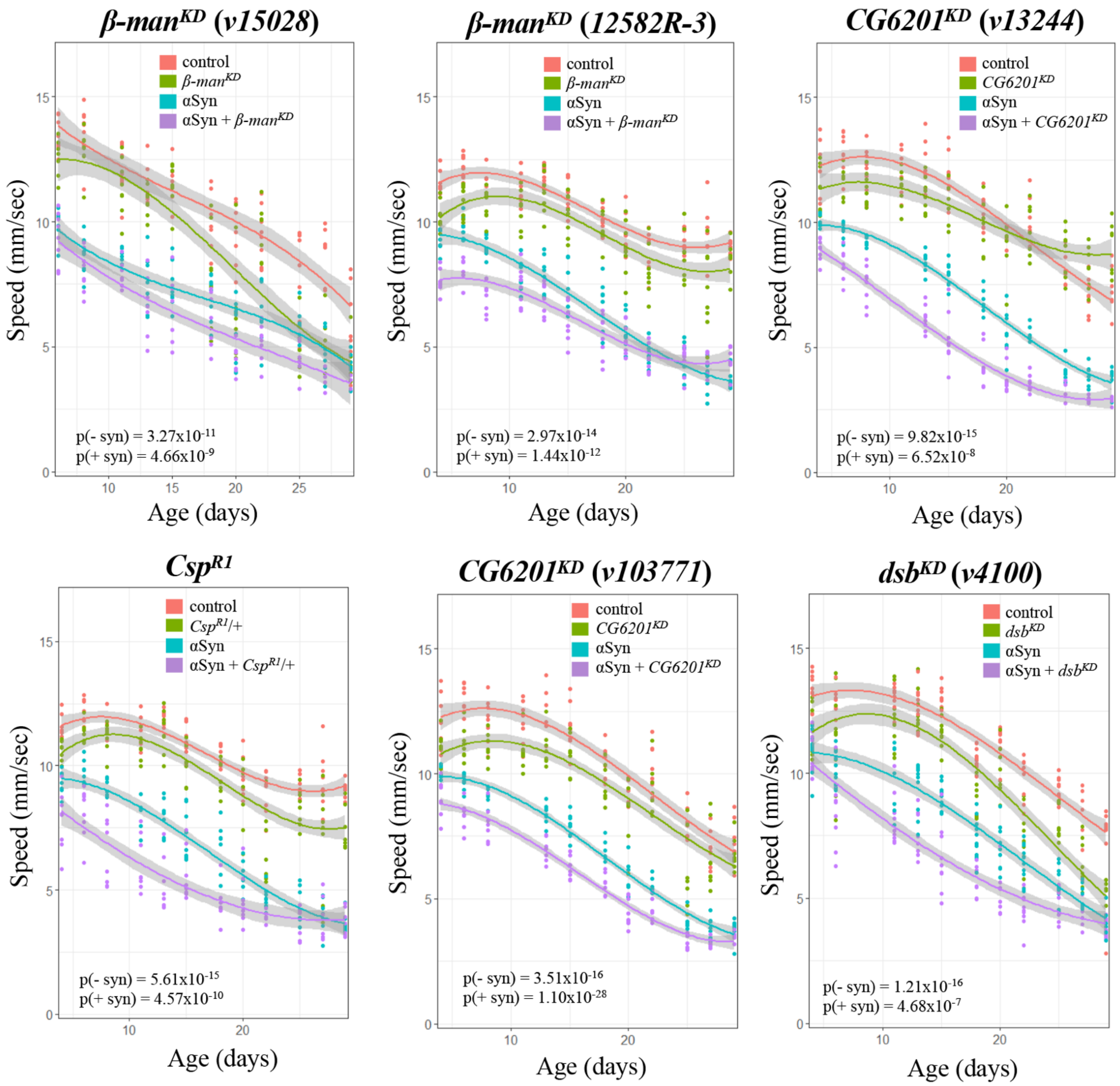
β-man^KD (v15028)
control
β-man^KD
αSyn
αSyn + β-man^KD
p(- syn) = 3.27x10^-11
p(+ syn) = 4.66x10^-9
Speed (mm/sec)
Age (days)
β-man^KD (12582R-3)
control
β-man^KD
αSyn
αSyn + β-man^KD
p(- syn) = 2.97x10^-14
p(+ syn) = 1.44x10^-12
Speed (mm/sec)
Age (days)
CG6201^KD (v13244)
control
CG6201^KD
αSyn
αSyn + CG6201^KD
p(- syn) = 9.82x10^-15
p(+ syn) = 6.52x10^-8
Speed (mm/sec)
Age (days)
Csp^R1
control
Csp^R1/+
αSyn
αSyn + Csp^R1/+
p(- syn) = 5.61x10^-15
p(+ syn) = 4.57x10^-10
Speed (mm/sec)
Age (days)
CG6201^KD (v103771)
control
CG6201^KD
αSyn
αSyn + CG6201^KD
p(- syn) = 3.51x10^-16
p(+ syn) = 1.10x10^-28
Speed (mm/sec)
Age (days)
dsb^KD (v4100)
control
dsb^KD
αSyn
αSyn + dsb^KD
p(- syn) = 1.21x10^-16
p(+ syn) = 4.68x10^-7
Speed (mm/sec)
Age (days)

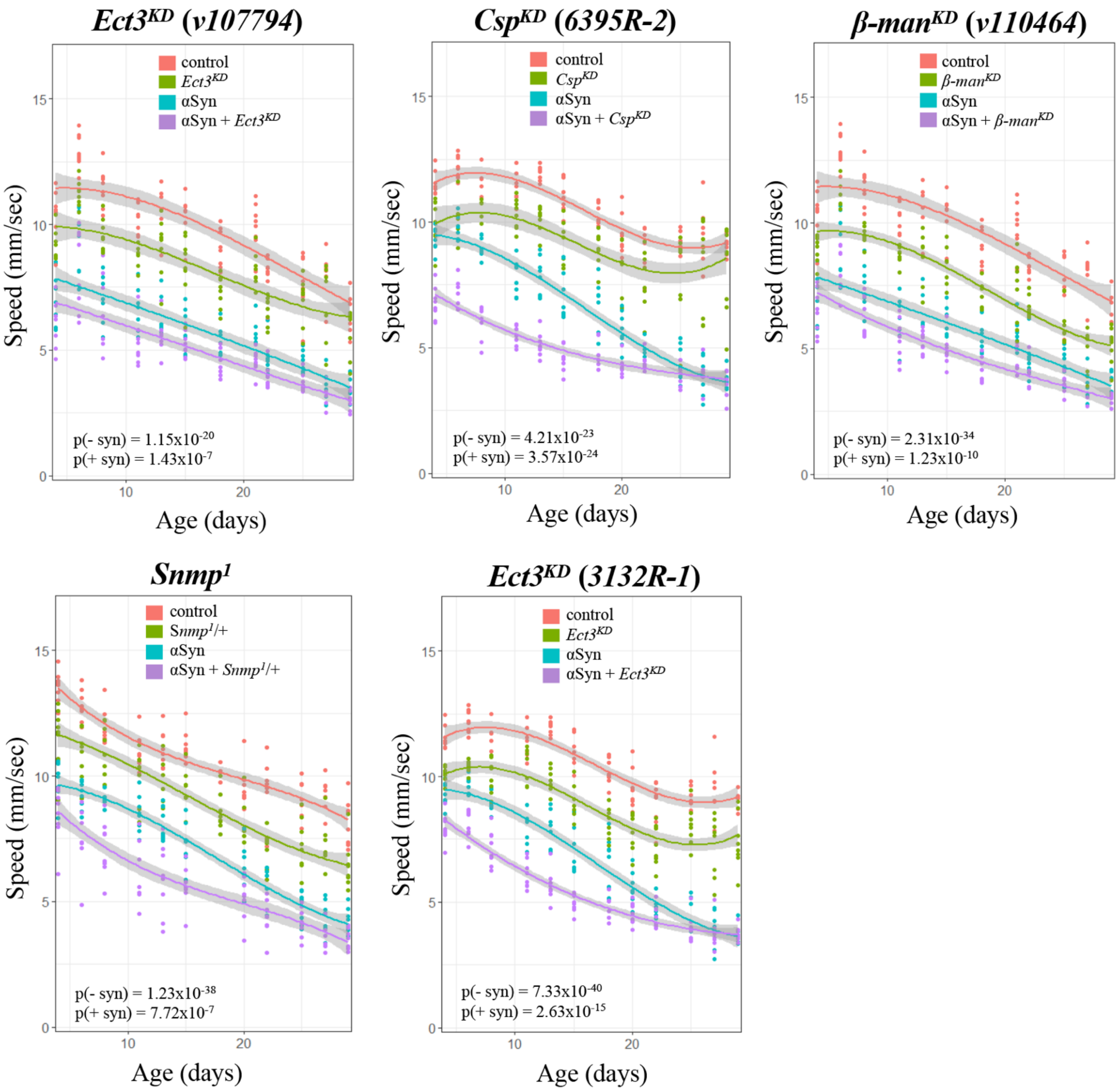
$Ect3^{KD}$ (v107794)
control
$Ect3^{KD}$
αSyn
αSyn + $Ect3^{KD}$
p(- syn) = $1.15x10^{-20}$
p(+ syn) = $1.43x10^{-7}$
Speed (mm/sec)
Age (days)
$Csp^{KD}$ (6395R-2)
control
$Csp^{KD}$
αSyn
αSyn + $Csp^{KD}$
p(- syn) = $4.21x10^{-23}$
p(+ syn) = $3.57x10^{-24}$
Speed (mm/sec)
Age (days)
β-$man^{KD}$ (v110464)
control
β-$man^{KD}$
αSyn
αSyn + β-$man^{KD}$
p(- syn) = $2.31x10^{-34}$
p(+ syn) = $1.23x10^{-10}$
Speed (mm/sec)
Age (days)
$Snmp^1$
control
$Snmp^1$/+
αSyn
αSyn + $Snmp^1$/+
p(- syn) = $1.23x10^{-38}$
p(+ syn) = $7.72x10^{-7}$
Speed (mm/sec)
Age (days)
$Ect3^{KD}$ (3132R-1)
control
$Ect3^{KD}$
αSyn
αSyn + $Ect3^{KD}$
p(- syn) = $7.33x10^{-40}$
p(+ syn) = $2.63x10^{-15}$
Speed (mm/sec)
Age (days)

**(C) severe toxicity independent of αSyn**

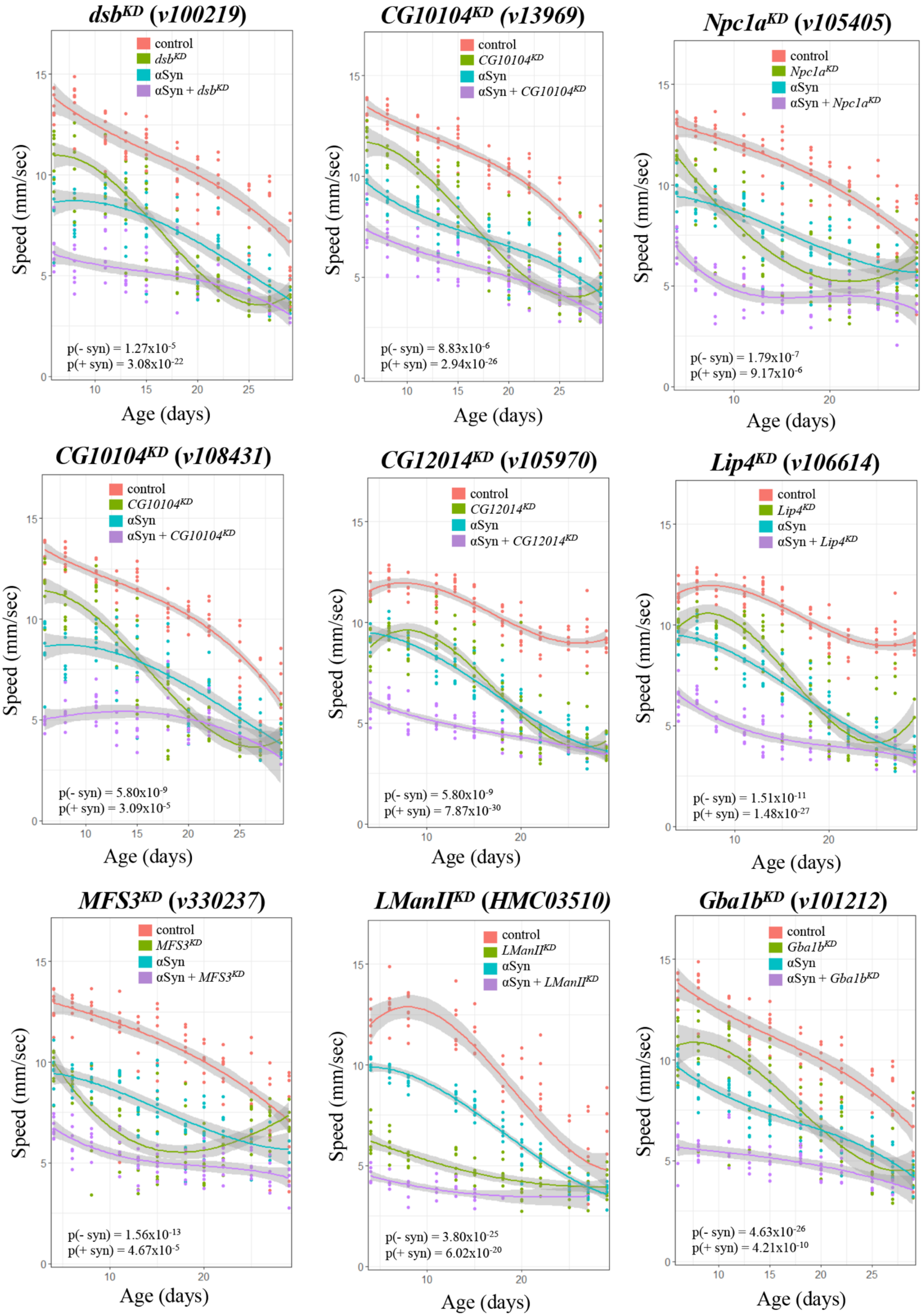

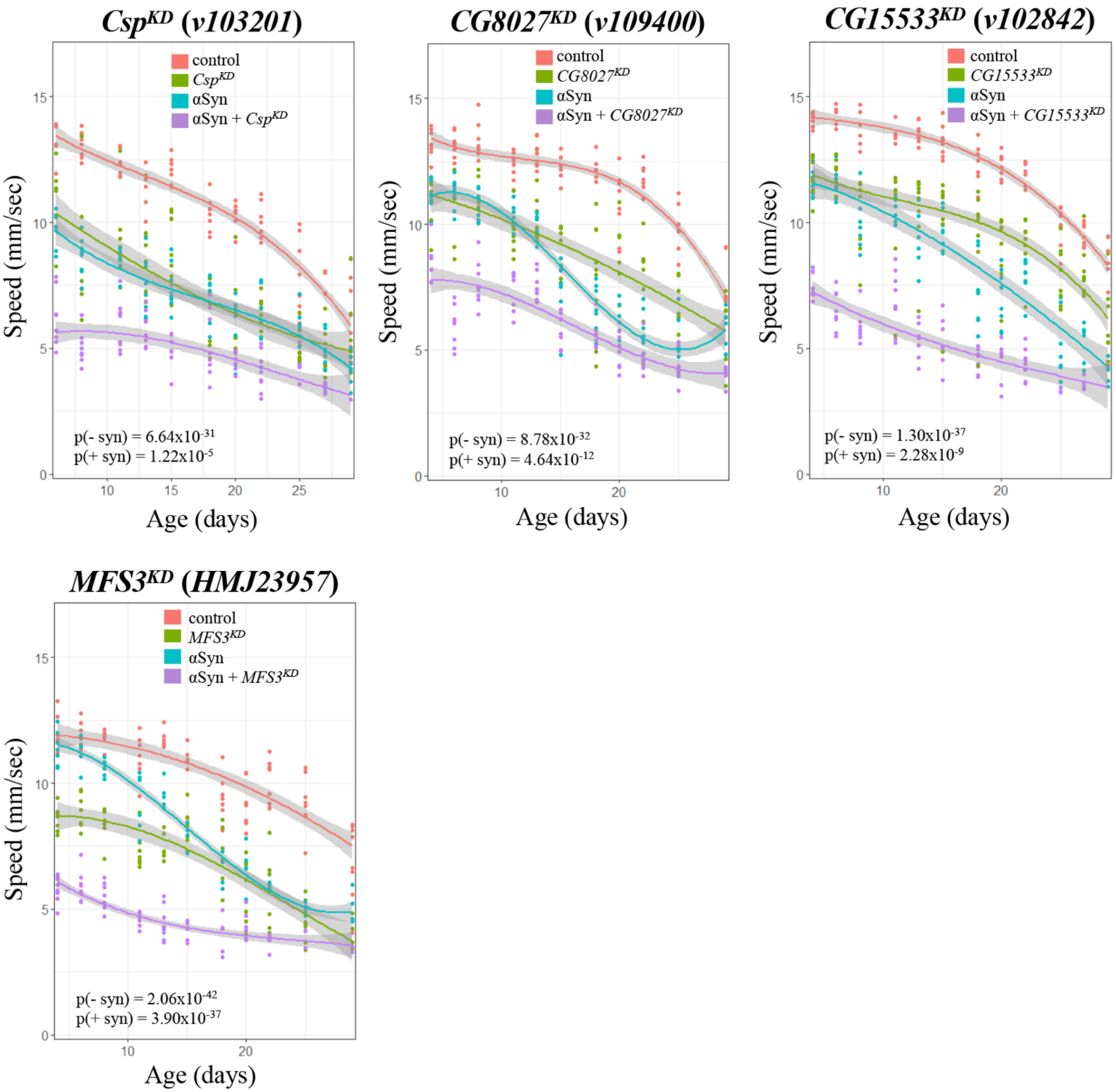

CspKD (v103201)
control
CspKD
αSyn
αSyn + CspKD
p(- syn) = 6.64x10-31
p(+ syn) = 1.22x10-5
Speed (mm/sec)
Age (days)
CG8027KD (v109400)
control
CG8027KD
αSyn
αSyn + CG8027KD
p(- syn) = 8.78x10-32
p(+ syn) = 4.64x10-12
Speed (mm/sec)
Age (days)
CG15533KD (v102842)
control
CG15533KD
αSyn
αSyn + CG15533KD
p(- syn) = 1.30x10-37
p(+ syn) = 2.28x10-9
Speed (mm/sec)
Age (days)
MFS3KD (HMJ23957)
control
MFS3KD
αSyn
αSyn + MFS3KD
p(- syn) = 2.06x10-42
p(+ syn) = 3.90x10-37
Speed (mm/sec)
Age (days)

**SUPPLEMENTAL FIGURE 2**

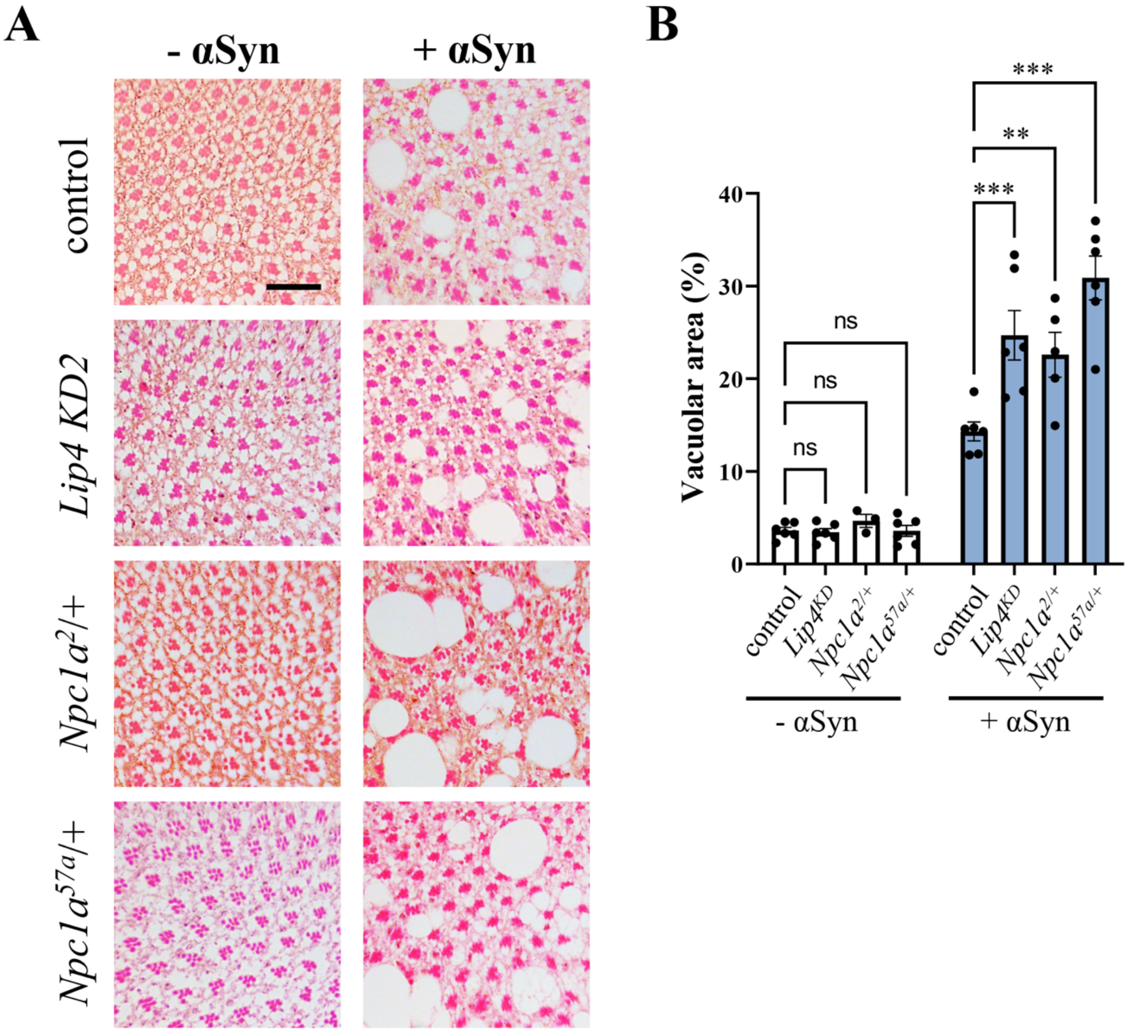


Representative images of retinal histology sections (A) and quantification (B) of additional *Npc1a* alleles and *Lip4* RNAi (*v106614*), showing consistent enhancement of α-synuclein-mediated neurodegeneration. Quantification based on extent of vacuolar changes (vacuole area / total area) from at least n=3 animals per genotype. Statistical comparisons were made using unpaired t-tests, followed by Dunnett's post-hoc test. Error bars represent the standard error of the mean. **, p<0.01; ***, p<0.001; ns, non-significant; Scale bar=20μm.
